# Genome Report: Diploid assembly of the Mexican lime genome reveals extensive heterozygosity

**DOI:** 10.1101/2024.09.30.615868

**Authors:** Isabelle Massaro, James Thomson, Aaron R. Leichty

## Abstract

Clonally propagated crops have long been recognized for their high levels of heterozygosity both between subgenomes within a somatic cell and between cells within an individual clone. Recent developments in long read sequencing technologies have accelerated our ability to identify this diversity and it is increasingly clear that these sources of diversity are abundant in clonal varieties and can contribute to variation in traits of interest to breeders. In this work, we assemble both subgenomes of Mexican lime (*Citrus* x *aurantifolia*), an interspecific hybrid between *C. hystrix* var. *micrantha* and *C. medica*. Using this chromosome-level assembly, we find extensive divergence between haplotypes, with at least 89% of the annotated genes harboring polymorphisms at an average rate of 13 per kilobase of coding sequence. Additionally, using high coverage PacBio HiFi libraries from leaf tissue of four individuals we identified multiple large structural variants differing between thorned and thornless lineages, and evidence for mosaicism at hundreds of loci. Many of these variants are found in the promoters and bodies of genes and may act as standing variation for continued improvement of this cultivar.

## INTRODUCTION

Perennial crops are often vegetatively derived selections from highly heterozygous ancestors. In most cases, these selections are propagated clonally, thereby maintaining the genetic diversity of the original parent. As a consequence, perennial crop systems often retain levels of genetic diversity closer to their wild ancestors than do many annual crops (Miller and Gross 2011). Historically, this retention of heterozygosity and high divergence between genome copies have slowed attempts to sequence and assemble perennial crop genomes. Longread sequencing technologies have overcome many of the hurdles limiting the simultaneous assembly of multiple haplotypes, including the need to generate highly inbred lines (e.g. grape genome, Jaillon et al. 2007) or elaborate methods to isolate homologous and homoeologous chromosomes (e.g. bread wheat, The International Wheat Genome Sequencing Consortium 2014). Within the past year alone, there have been at least 50 reports of new haplotype resolved plant genomes (NCBI Pubmed search), with the majority from woody or perennial systems (e.g. Yocca et al. 2024; Miao et al. 2024; Wang et al. 2024).

Previous work in other clonally propagated systems have revealed the necessity of accounting for heterozygosity when targeting genes via CRISPR. For example in the hybrid poplar clone 717 (*Populus tremula* x *P. alba*) it was demonstrated that polymorphism between alleles resulted in unequal editing of target genes (Zhou et al. 2015). Additionally unaccounted for halplotype differences significantly reduces mapping quality in RNA-seq, DNA-seq and ChIP-seq experiments (Xue et al. 2015; Shi et al. 2024). Conversely, in clonal systems where the CRISPR editing machinery cannot be removed, and editing often results in biallelic loss-of-function mutations, heterozygosity may give researchers added control over the degree of gene disruption. Together, these findings have been a strong motivation to generate genome assemblies that account for these differences between haplotypes (Xue et al. 2015; R. Zhou et al. 2023).

In addition to their standing diversity between haplotypes, clonal systems are increasingly appreciated for their high degree of mosaicism. Recent work in clementine trees estimated that a mature tree has 1,500-5000 somatic mutations, and that ∼1 new mutation arises with each flush of new leaves (Perez-Roman et al. 2022). Since citrus is often clonally propagated through apomictic seed, some portion of these mutations will be carried through each clonal generation. This mosaicism is often a target of selection for variation in pigmentation of leaves, flowers, and fruits, and seems to contribute to trait improvement even when phenotypes are less readily visible (Ohtsu and Kuhara 1994). Interestingly, there are now multiple examples to suggest this variation is a powerful source of standing variation in clonal systems. For example, in cassava, it appears resistance to Cassava mosaic disease is mediated by chimerism of a polymorphism in a *DNA polymerase* subunit and is maintained as such in much of the clonal cassava germplasm (Lim et al. 2022). Recent work in sweet orange found that variants fixed between different selections (somatic mutations) explain the differences in fruit size between varieties (Wang et al. 2024). Ultimately, these mutations would have been selected from chimeric plants. With the advent of haplotype-resolved assemblies it should be possible to identify somatic variants available for selection and may ultimately accelerate trait improvement.

To this end, we report the construction of a chromosome-scale and haplotype-resolved assembly of the Mexican lime genome (*Citrus* x *aurantifolia*) an interspecific hybrid of a micrantha (*C. hystrix* var. *micrantha*) and a citron (*C. medica*) (Wu et al. 2018). Mexican lime, also known as Key lime or bartender’s lime is valued its juice that is more acidic and aromatic then it’s larger relative the Persian or Bearss lime (Figure 1). The plant is generally shrub-like, but can grow to over 15 feet on its own rootstock. Relative to other varieties of citrus, Mexican lime has few developed varieties, yet using the new genome and long read sequencing of multiple thorned and thornless varieties we identify many large structural variants and mosaic loci. These loci may be a novel source of variation in breeding programs for future development of this variety.

**Figure 1.**
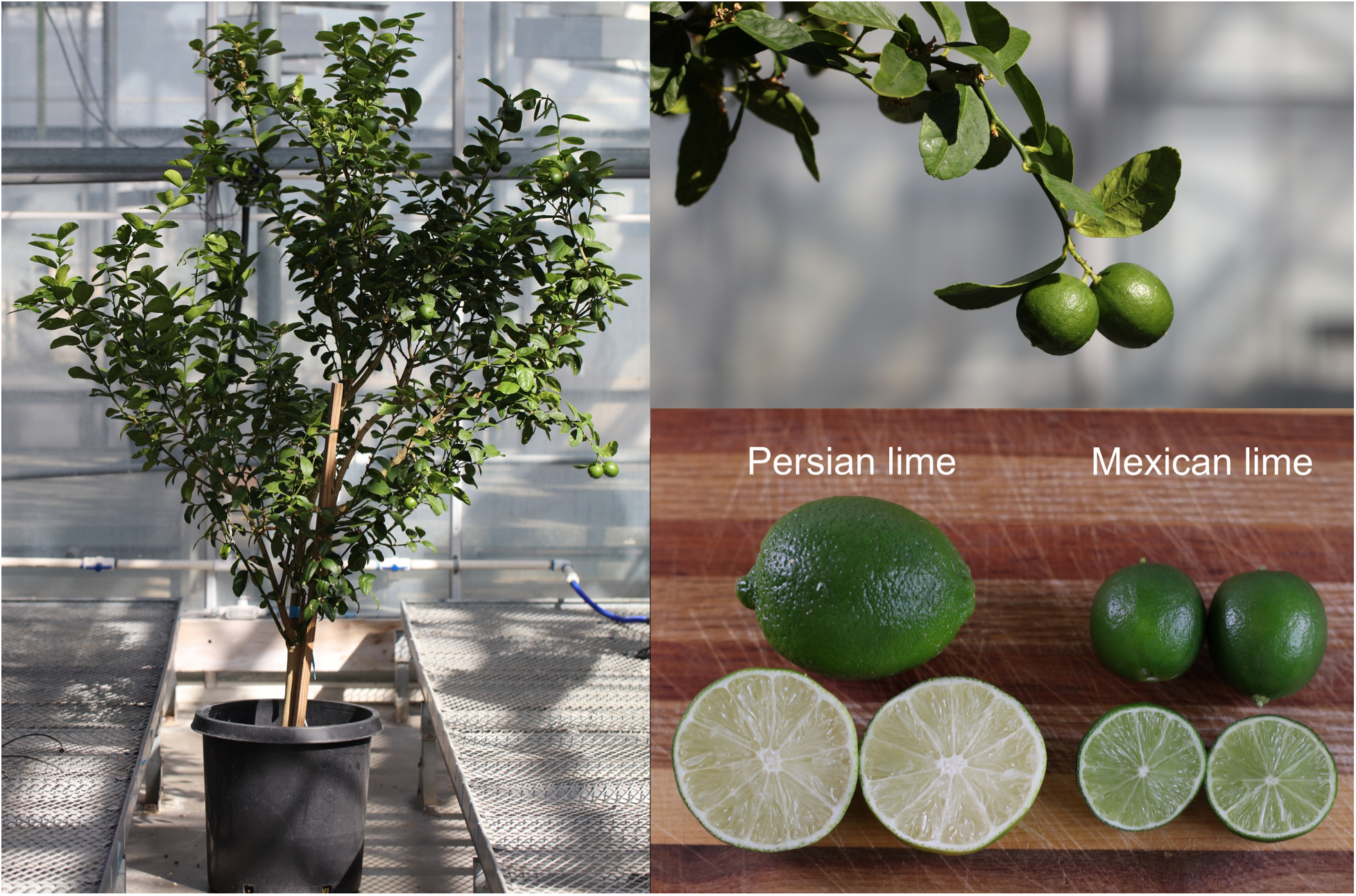
Mexican lime tree and fruit compared with Persian lime.

## METHODS & MATERIALS

### Plant material

For genome assembly of Mexican lime, leaf tissue from a mature thornless tree (TLR) was used (purchased from East Bay Nursery, Berkeley, CA). Additionally, leaves from a second thornless tree (TL1) and two thorned trees (TD1 and TD2) were used for variant analysis (each was obtained from a unique commercial source).

For RNA-seq, trees were grown in a greenhouse under 16 hrs. light/ 8 hrs. dark and grown in 10 gallon pots with regular fertilization.

### PacBio Library and Sequencing

HMW DNA was extracted from young leaves and shoot apices of TLR (BioSample SAMN42593883) using an in-house CTAB protocol at Brigham Young University DNA Sequencing Center (Provo, UT). The PacBio SMRTbell library (15-20kb) was constructed using SMRTbell Express Template Prep Kit 3.0 (PacBio, Menlo Park, CA, USA) using the manufacturer recommended protocol. The library was loaded onto a 8M SMRT cell and run on a PacBio Revio for 24 hours. The three additional samples not used for assembly were also sequenced in the same manner (TL1, TD1, and TD2)

### HiC library preparation and sequencing

Nuclei were isolated from fresh-frozen plant tissue samples using the CelLyticTM Plant Nuclei Extraction Kit (Sigma). Hi-C data was generated utilizing the High Coverage HiC kit from Arima, adhering to the manufacturer’s instructions. In brief, chromatin was cross-linked with formaldehyde and digested with a proprietary cocktail of restriction enzymes. Following this step, the 5’-overhangs were filled in and labeled with a biotinylated nucleotide and spatially proximal digested ends of DNA were ligated. The ligated DNA was then purified and sheared to 550–600 bp by sonication and the biotinylated fragments were selectively enriched. Subsequently, the enriched proximally-ligated DNA fragments were subjected to library preparation using KAPA HyperPlus Kits (Roche), including end-repair, dA-tailing, adapter ligation and PCR amplification. Finally, the Hi-C library was sequenced on the Illumina Novaseq X plus platform with 150 bp paired-end reads. Raw reads (Mxlime_HiC; Table S1) were processed using the ArimaGenomics mapping pipeline (https://github.com/ArimaGenomics/mapping_pipeline).

### mRNA-seq of various plant tissues

Mature leaves, flowers, young fruit, and vegetative shoot apices (Figure S1) from the same plant used for genome assembly (BioSample SAMN42593883) was used for RNA extraction with the Qiagen Rneasy Plant Mini Kit using the Sigma On-column DNase digestion kit. The NEBNext Ultra II Directional RNA Library Prep Kit was used to construct PolyA libraries according to the manufacture’s protocol (New England Biolabs). The libraries were sequenced with PE150 format by Admera Health (South Plainfield, New Jersey).

### Genome assembly

PacBio HiFi reads were assembled using hifiasm (v0.19.5) using HiC reads to generate a haplotype-resolved assembly (Cheng et al. 2021; 2022). Contigs from both haplotypes over 1 Mb in size were then used for scaffolding using SALSA with the same HiC reads (Ghurye et al. 2019; 2017). Manual corrects were made to the resulting scaffolds, including two breaks and multiple joins to obtain the final pseudomolecule assembly. All chromosomes were oriented in the same direction as *C. sinensis* DHSOv3 (Wang et al. 2021). In the case where scaffolds were manually joined, whole genome alignments with the DHSOv3 assembly and the *C. australis* v1.0 (UQ) assembly (Nakandala et al. 2023) were used to determine pairing.

### Gene prediction and annotation

*De novo* identification and classification of repeats was done using RepeatModeler v.2.0.1 (Flynn et al. 2020). RepeatMasker v.4.0.7 (Smit, Hubley, and Green 2013) was used with the Mexican lime repeat library to soft and hardmask the genome using the following parameters: -nolow -norna -xsmall.

BRAKER3 was used to annotate genes on the softmasked genome (Gabriel et al. 2023; Hoff et al. 2016; Brůna et al. 2021; Hoff et al. 2019). First, fastp was used for trimming and quality filtering of RNA-seq reads (Chen et al. 2018). Hisat2 v2.2.1 (Kim et al. 2019) was used to map the RNA-seq libraries and the resulting alignments were supplied to the BRAKER pipeline along with UniProt proteins from the Viridiplantae dataset (The UniProt Consortium 2021). Additionally, StringTie (Pertea et al. 2015) is used to assemble the RNA-seq reads followed by rounds of GeneMark and AUGUSTUS training and gene prediction (Stanke and Waack 2003). Finally, gene sets were combined with TSEBRA (Gabriel et al. 2021).

### Haplotype phasing and construction of pseudomolecules

SubPhaser was used to determine the chromosome sets from each parent species (*C. medica* and *C. hystrix*) (Jia et al. 2022). Following subgenome assignment, Mashmap (Jain et al. 2018) was used to identify the *C. medica* subgenome by aligning chromosomes with the *C. medica* RLv1 assembly (Wang et al. 2017).

### Analysis of nucleotide and structural variants

SNPs and indels were determined using Minimap2 (v2.28-r1209) to generate alignments between haplotypes (Li 2018; 2021). For this, the hystrix haplotype was set as reference and the “asm20” option was used to allow for up to 20% divergence. Bedtools (v2.27.1) was used to determine the feature location of variants (Quinlan and Hall 2010).

The identification of population variants was done using PacBio HiFi libraries of leaf tissue from three individuals (1 thornless, and 2 thorned) as well as a random subset of the original reads used for generation of the Mexican lime assembly (a thornless individual). Reads were aligned to the diploid assembly using nglmr v0.2.7 (Sedlazeck et al. 2018) and variant calling was done using Sniffles2 v2.2 (Smolka et al. 2024). SNPs were also called from these same alignments using BCFtools v1.21 (Danecek et al. 2021). Read alignments and variant calls were visualized using Jbrowes version 2 (Diesh et al. 2023).

### Genome assessment and comparative genomics

Assessment of genome completeness was conducted with BUSCO v5 using the Embryophyta *odb10* dataset (Manni et al. 2021). Alignment of DNA-seq reads was done with bowtie2 v2.4.4 using the following options: --end-to-end –sensitive (Langmead and Salzberg 2012). Alignment of RNA-seq reads were done with Hisat2 v2.2.1 (Kim et al. 2019).

Alignment of the hardmasked Mexican lime genome with the *C. sinensis* di-haploid genome DHSOv3 (Wang et al. 2021) was done using MashMap (Jain et al. 2018) and the following parameters: -f none --pi 98. Macrosynteny analyses were done using the MCscan pipeline (Tang et al. 2008). The details of these analyses can be found at the DRYAD submission associated with this paper.

## RESULTS AND DISCUSSION

### Construction of phased pseudochromosomes

*De novo* assemblies of both Mexican lime haplotypes were constructed using 46.5 Gb of PacBio HiFi reads with a median read length of 12,620 bp. This represents approximately 60X coverage of each haplotype assuming a haploid genome size of 392 Mb (Ollitrault et al. 1994). The initial haplotype assemblies were then scaffolded using HiC conformation capture reads to generate a final diploid assembly of 18 chromosomes and 8 unplaced scaffolds ranging from 1.8 to 7.5 Mb in size. The total size of the assembly was 708 Mb, representing 90% of the predicted genome size (Table 1).

**Table 1.** Statistics for the *C. x aurantiifolia* pseudomolecule assemblies.

| Species | *hystrix*<br>haplotype | *medica*<br>haplotype | *C. sinensis* doubled haploid*<br>(Wang et al. 2021) |
| --- | --- | --- | --- |
| Chromosome<br>number (n) | 9 | 9 | 9 |
| Assembly size<br>(Mb) | 325 | 355 | 304 |
| Number of<br>predicted<br>protein coding<br>genes | 23,936 | 24,334 | 29,268 |
| Busco genome<br>(embryophyta)** | C:98.4%<br>S:97.3%<br>D:1.1% | C:98.6%<br>S:97.5%<br>D:1.1% | C:98.3%<br>S:97.3%<br>D:1.0% |
| Busco protein<br>(embryophyta)** | C:98.4%<br>S:79.8%<br>D:18.6% | C:97.9%<br>S:79.6%<br>D:18.3% | C:96.6%<br>S:58.6%<br>D:38.0% |

\*Analysis based on the 9 chromosomes only (excluding chrUn and associated genes).
\*\*percent of embryophyta gene models or proteins that are present as complete (C), and portion that exist as single-copy (S) or duplicates (D) in the assembly.

To determine the subgenome origin of each chromosome we utilized SubPhaser (Jia et al. 2022) to identify subgenome specific repeats. This analysis identified 8,388 repeats (15-mers) that were differentially enriched between chromosome pairs. These repeats were used for clustering based on their frequency across each chromosome (Figure S2) leading to unambiguous assignment of each chromosome haplotype to either subgenome (Figure 2 and S3). The distribution of haplotype enriched long terminal repeat retrotransposons (LTR-RTs) mirrored that of the kmers (Figure 2). Based on this analysis there were some evidence of exchange and/or assembly errors between homoeologous chromosomes (e.g. right arm of chr4_medica; Figure 2, rings 2 and 3), but the majority of these blocks did not extend for more than 1 Mb. In general, and consistent with what is known about Mexican lime ancestry (Wu et al. 2018), the subgenomes show minimal evidence for exchange compared with other recently analyzed interspecific hybrids like *Populus alba* x *Populus tremula* (Jia et al. 2022), or *Citrus sinensis* (N. Wang et al. 2024).

**Figure 2.**
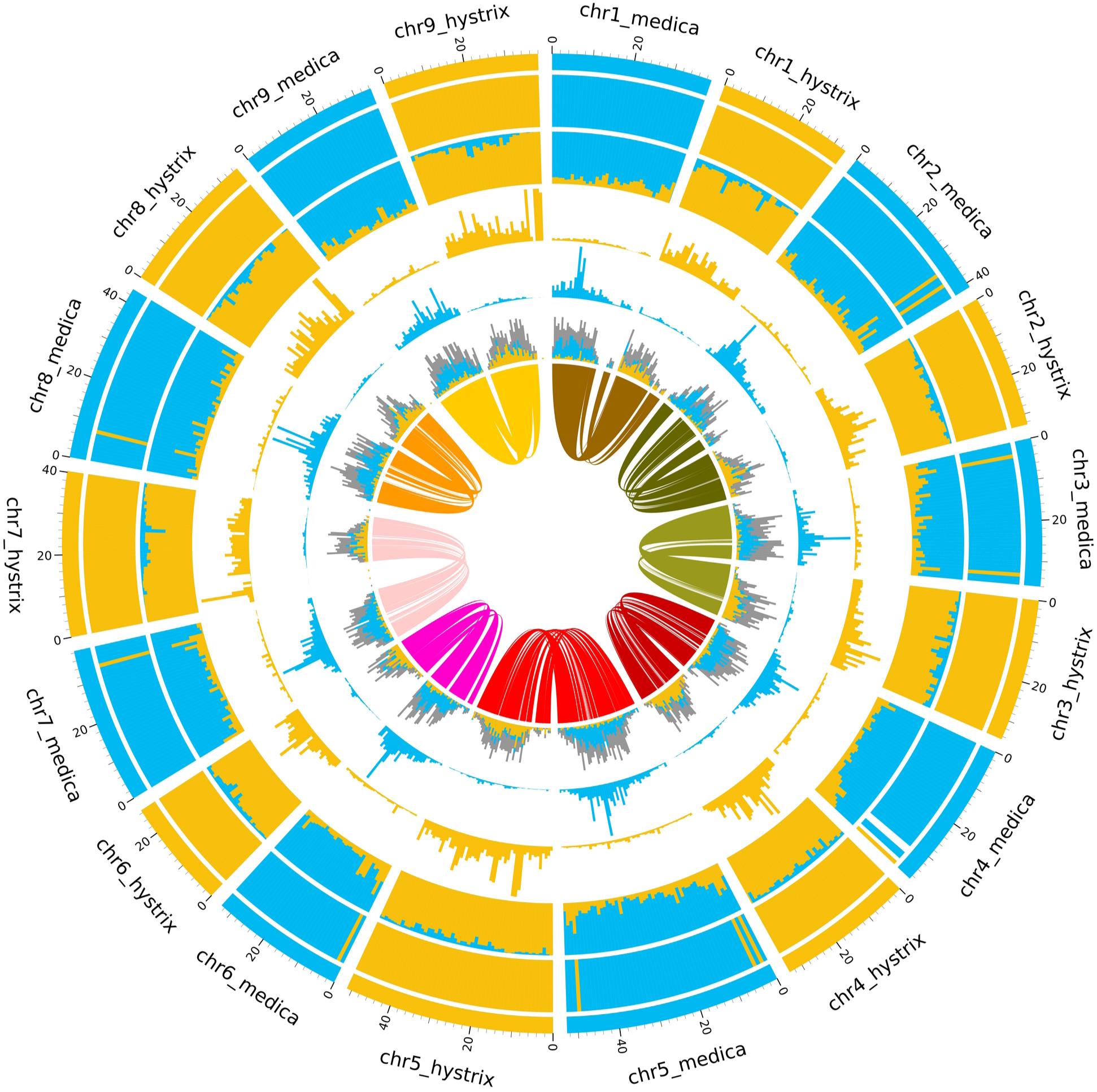
Subgenome assignment of chromosomes. Starting from the outer ring of the circos plot (1-7): (1) subgenome assignment of each chromosome using k-means clustering (either yellow or blue); (2) enrichment of subgenome-specific kmers for 1 Mb sliding windows across each chromosome; white areas represent no significant enrichment of either subgenome; (3) proportion of subgenome-specific k-mers for each interval; (4-5) counts for subgenome-specific k-mer sets; (6) enrichment of long terminal repeat retrotransposons (LTR-RTs) across each chromosome interval; color represents significant enrichment for a specific subgenome, whereas grey represents non-specific LTR-RTs; (7) homeologous blocks across genome.

### Assessment of genome completeness

To annotate the haplotype assemblies, we utilized the BRAKER3 pipeline and a set of mRNA-seq libraries from aerial tissues of mature plants (Figure S1). In total, we identified 23,936 and 24,334 protein coding genes for the hystrix and medica haplotypes, respectively (Table 1). By comparison, a recent assembly of *C. sinensis* DHSOv3 (Wang et al. 2021) found 29,268 protein coding genes in their doubled haploid genome. However, BUSCO analysis of the Mexican lime haplotype sequences found comparative levels of completeness to the DHSOv3 assembly. Interestingly, BUSCO analysis on the protein sets of each haplotype showed higher levels of completeness for the Mexican lime assembly (Table 1, hystrix: 98.4% and medica: 97.9% versus DHSOv3: 96.6%). Incorporation of more extensive RNA-seq datasets spanning more tissue types and conditions will likely even the numbers of protein coding genes between Mexican lime and other cultivars.

Comparison of each haplotype to the DHSOv3 assembly shows end-to-end representation for all chromosomes (Figure 3). Indeed, 4 of 9 and 6 of 9 chromosomes from the hystrix and medica haplotypes, respectively, contained telomeres at both ends (Table S2). Additionally, all of the remaining chromosomes harbored a telomere at one end. Interestingly, there were minimal structural differences between haplotypes and the DHSOv3 genome, with 3 inversions on chromosomes 2, 7, and 8, and 1 translocation on chromosome 5. Additionally, there was no evidence for interchromosomal translocation or duplication (Figure 3).

**Figure 3.**
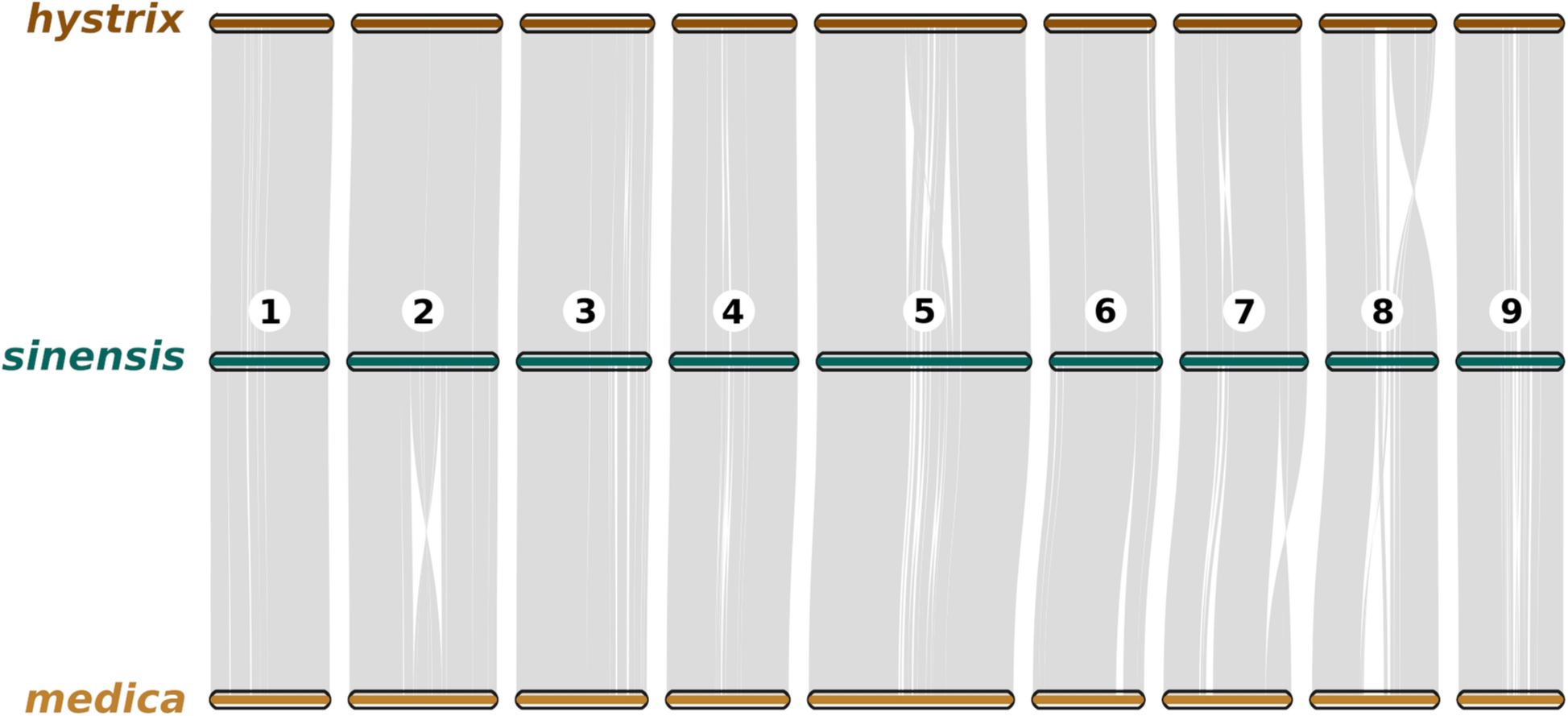
Comparison of each Mexican lime haplotype to the *C. sinensis* DHSOv3 genome (Wang et al. 2021).

### Comparison of haplotypes and characterization of structural variants

To quantify difference between haplotypes we utilized whole genome alignment in Minimap2, consistent with methods used by other recently published genomes (e.g. (Wang et al. 2024). Using this method, we were able to only align 64 Mb between the two haplotypes, whereas the same technique yielded 119 Mb between sweet orange haplotypes. Given that Mexican limes are known to harbor some of the highest heterozygosity of citrus varieties (Wu et al. 2018), we also compared genes in syntenic blocks between haplotypes. Of the 21,518 syntenic pairs (∼90% of all annotated genes), there was an average rate of 13 SNPs or InDels per 1 kb of coding sequence. In total, only 154 genes had identical coding sequence between haplotypes. Given that our annotation does not include untranslated regions, it’s likely that nearly every gene in the genome is distinguished by at least one polymorphism.

Given that Mexican lime clonally propagated from seed we reasoned that there should be high levels of mosaicism within the tissue of individuals. To allow for the identification of such potential variation we sequenced an additional three individuals, two thorned and one thornless, to high depth using PacBio HiFi reads (20-30X). Additionally, we remapped the reads from the thornless individual used for genome assembly. Together across all four samples, we found 374 structural variants larger than 50 bp (Figure S4). Of the different variant types, deletions and insertions were most frequent (149 and 123, respectively). We also found evidence for 61 breakend events, 39 duplications and 2 inversions. Of these variants the majority were either present in only the two thorned individuals (n=141, Figure S5) or present in all 4 individuals (n=76). There were also a number of variants specific to either thorned plant (n= 63 and n=38). For the variants shared between thorned plants, the majority were present as heterozygous genotype calls (Figure S6). Given that we mapped our reads to the diploid assembly, this suggest that these variants are the result of duplications of loci present as single loci in the thornless genotypes or these loci are chimeric in the thorned plants (Figure S7 and S8).

Using the HiFi reads, we also attempted to identify chimeric SNPs. Using a standard SNP calling pipeline, we identified over 33,000 variants across all 4 plants, of which ∼5,000 had a heterozygous genotype in thornless plants. However, thorned plants had more than 20,000 sites with a heterozygous genotype. This disparity suggests that there are a number of large duplications in the thorned genome (see Figure S8) that results in allele frequency depths consistent with a heterozygous genotype (instead of homozygous at two copies of a region). However, this possibility would not explain the putatively mosaic sites in the thornless plant for which the genome was assembled. Indeed, the majority of these heterozygous positions appear to be real mosaic loci in the leaf tissue that was sampled (Figure S9). Future work attempting an independent assembly of a thorned genotype would help to resolve these possibilities, as well as deeper short read sequencing of a more diverse panel of Mexican lime lineages. Together these analyses suggest substantial standing variation between haplotypes and within tissues that breeding efforts could leverage in the future.

## DATA AVAILABILITY

The genome assembly and raw sequencing data have been submitted to Genbank under the bioproject: PRJNA1137419. A breakdown of the associated SRA accessions can be found in Table S2. Genome annotations and additional assembly data can be found on DRYAD.

## ACKNOWLEDGMENTS

We are thankful to the staff at Admera Health, Quick Biology, and the BYU DNA Sequencing Center for help obtaining the sequencing data.

---

**Figure S1.**
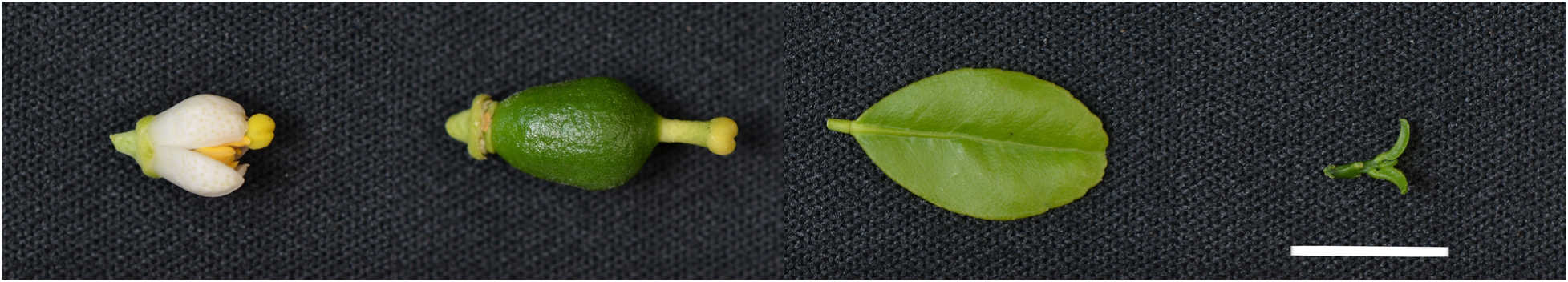
Samples used for RNA-seq analysis. From left: flower, fruit, leaf, apex. Scale bar is 1 cm.

**Figure S2.**
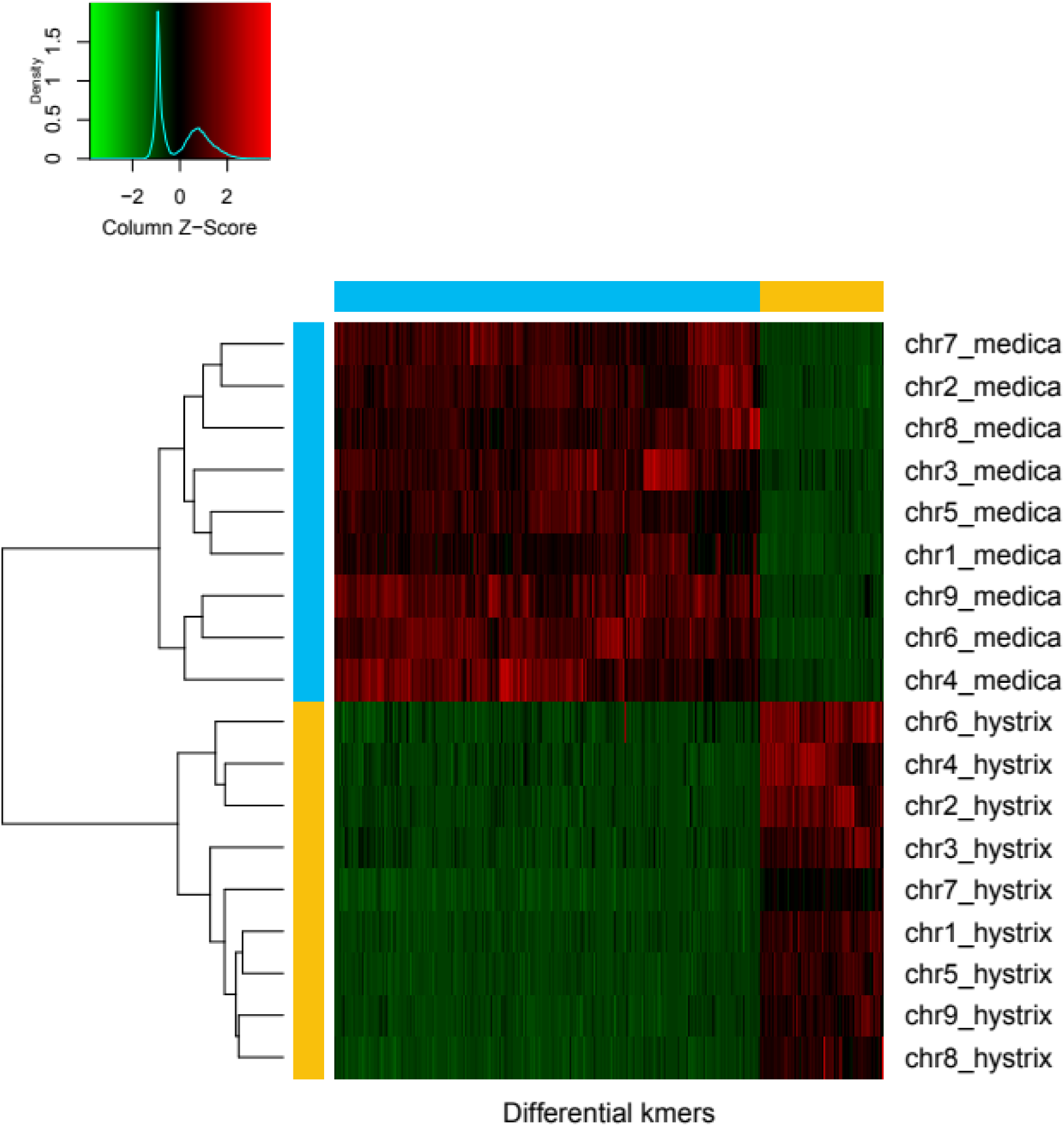
Hierarchical clustering of 15-mers enriched in each subgenome. Blue and yellow bars denote subgenome assignment of chromosomes listed on the right side of the figure. Coloring of the heat map for each unique 15-mer corresponds to a releative Z score, with red denoting high enrichment in either the blue or yellow subgenomes.

**Figure S3.**
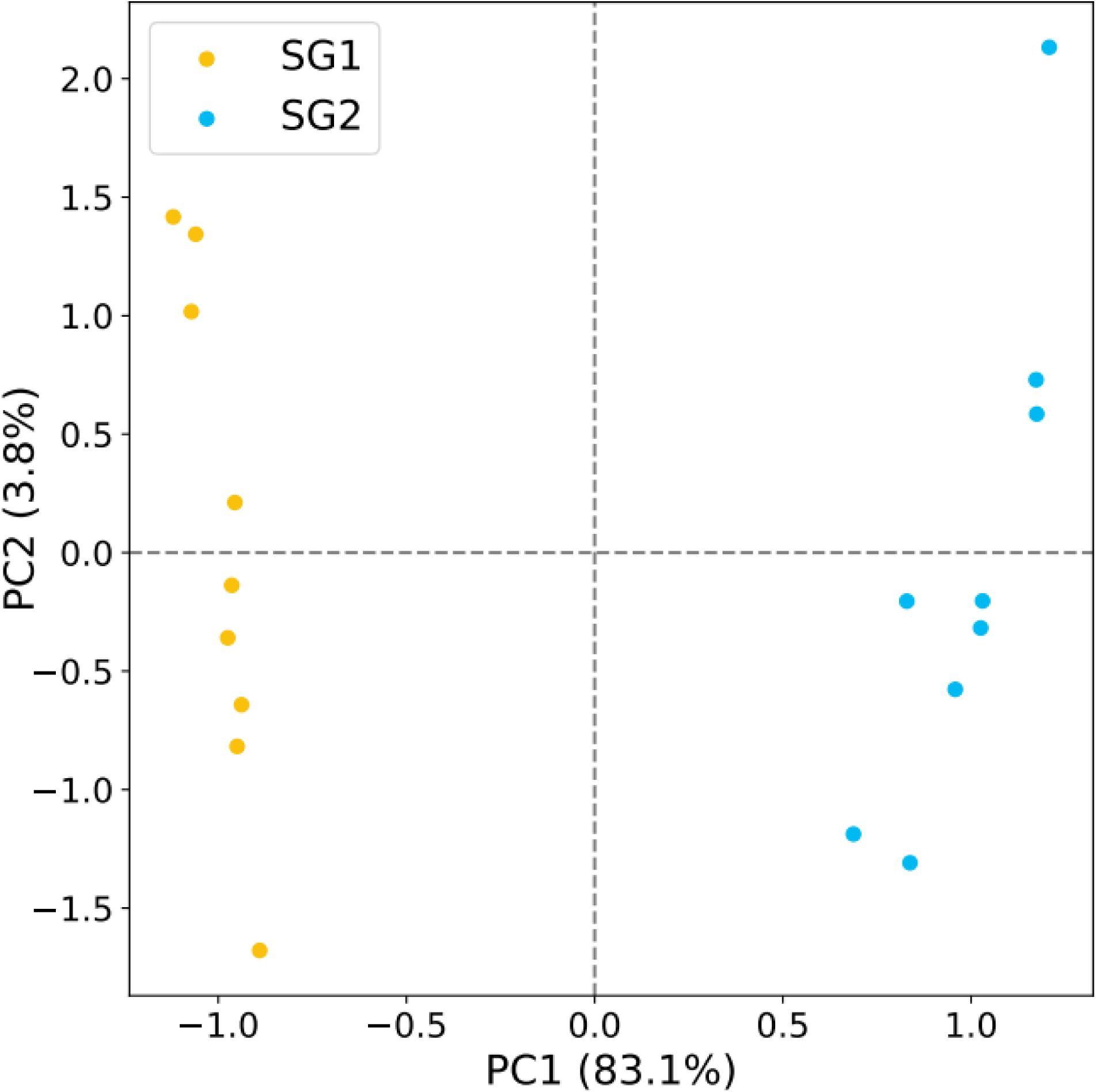
Principle component analysis (PCA) of the subgenome-specific 15 mers.

**Figure S4.**
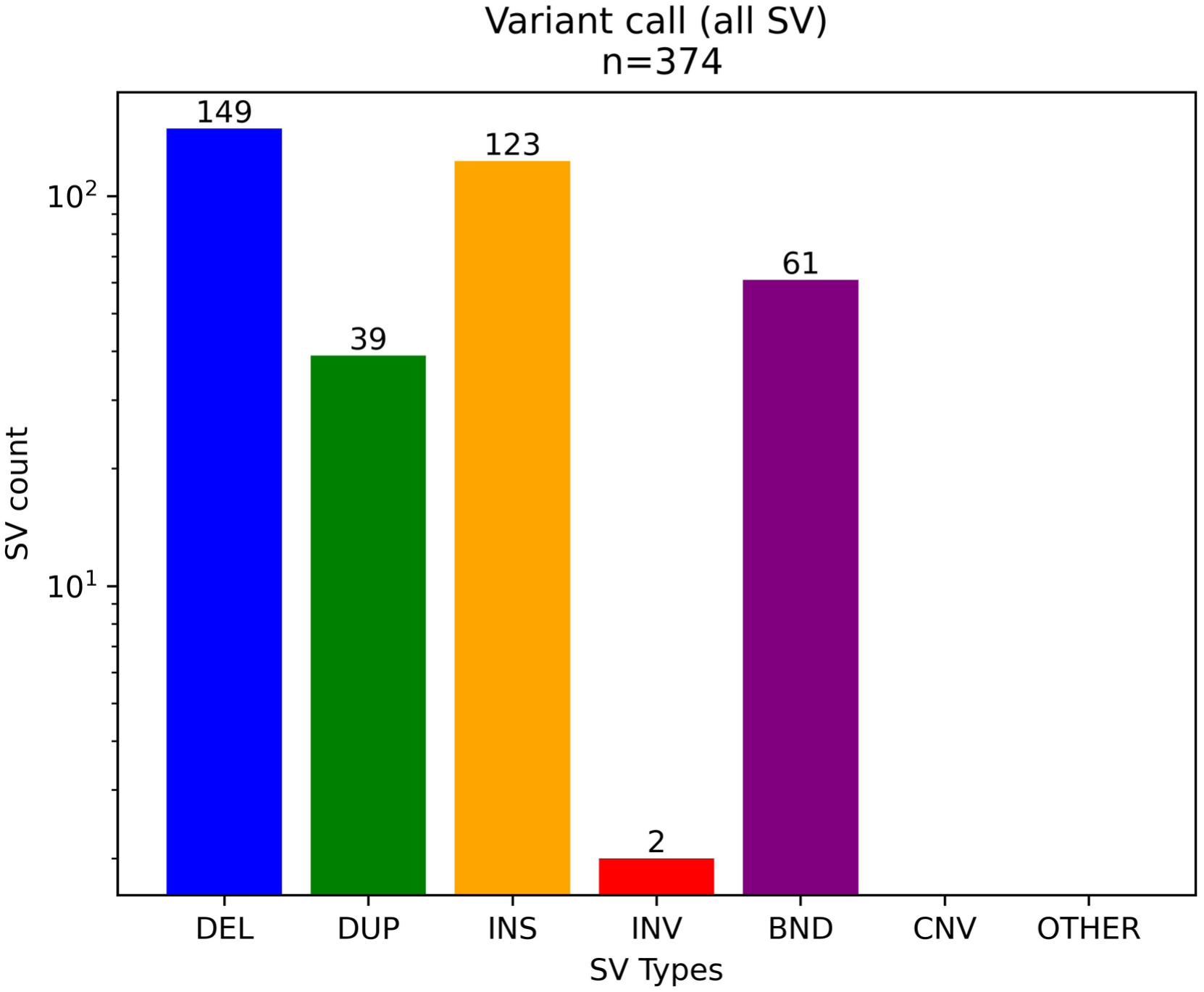
Counts for Sniffles2 structural variant calls. DEL = deletion. DUP = duplication, INS = insertion, INV = inversion, BND = breakend event.

**Figure S5.**
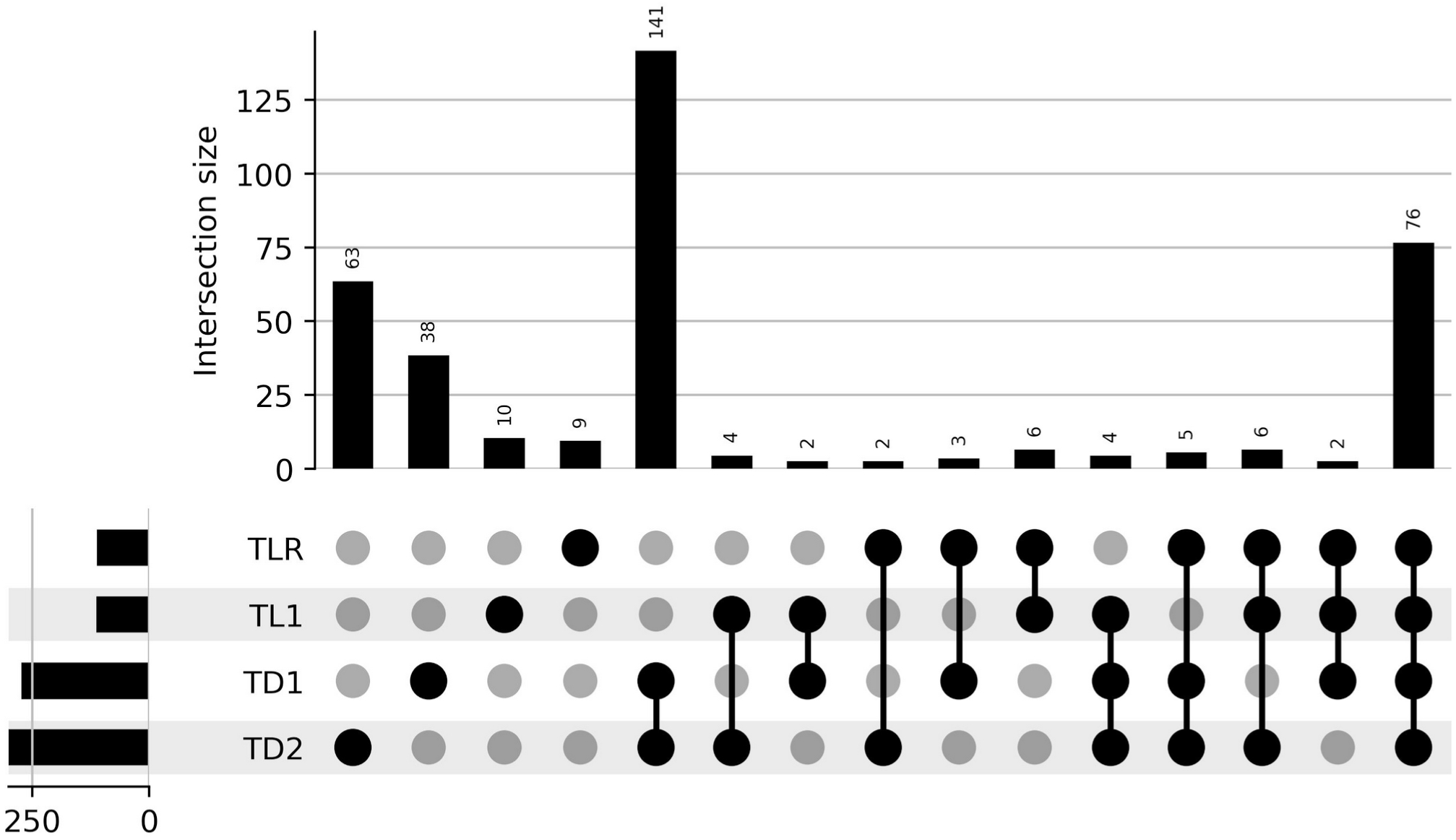
Shared and unique variants identified using Sniffles2 and the PacBio HiFi datasets from two thornless (TLR and TL1) and two thorned individuals (TD1 and TD2) relative to the diploid Mxlime_USDA_v1 assembly. Only variants of 50 bp or larger were considered. Variants are considered shared if present as either a 0/1 or 1/1 genotype in each individual.

**Figure S6.**
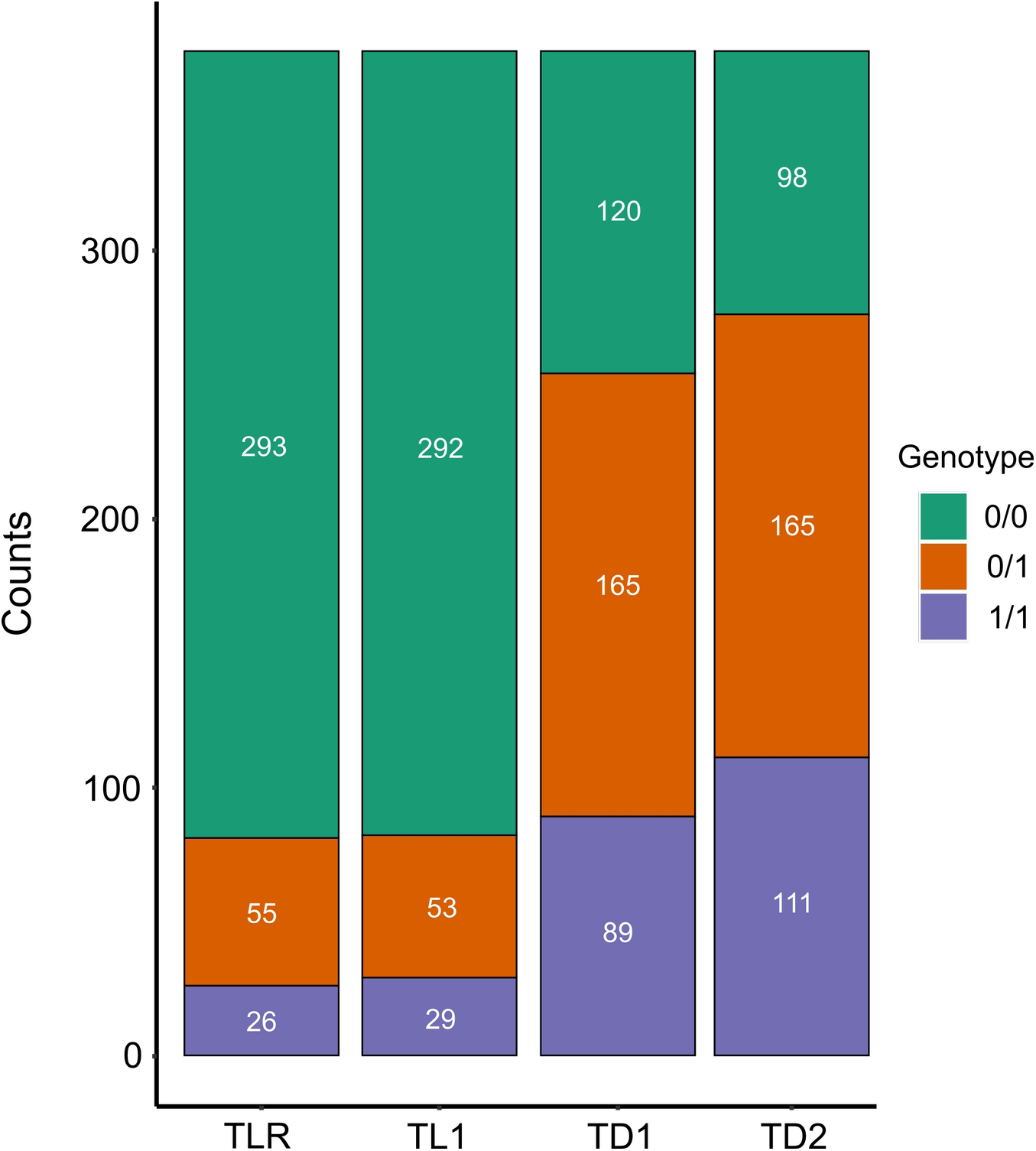
Genotype counts for structural variants identified using PacBio HiFi datasets from two thornless (TLR and TL1) and two thorned individuals (TD1 and TD2) relative to the diploid Mxlime_USDA_v1 assembly. Only variants of 20 bp or larger were considered. Variants with a 0/1 genotype are potential mosaic variants found in the leaf tissue sampled for the PacBio libraries.

**Figure S7.**
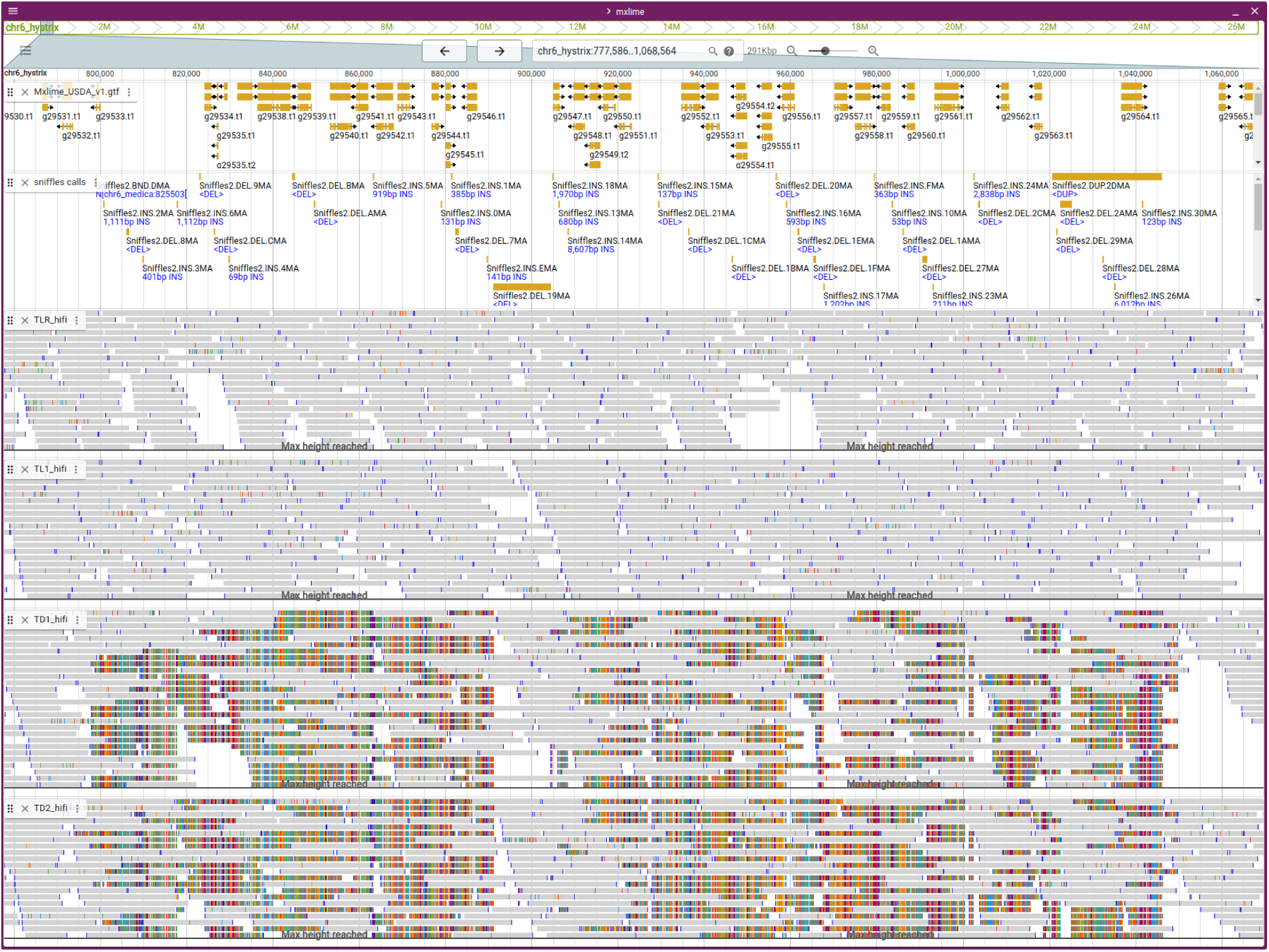
JBrowse2 view of putative large duplication event unique to thorned individuals. Duplication was not called by Sniffles2, but is instead associated with many small structural events in the region (i.e. 2^nd^ track from the top).

**Figure S8.**
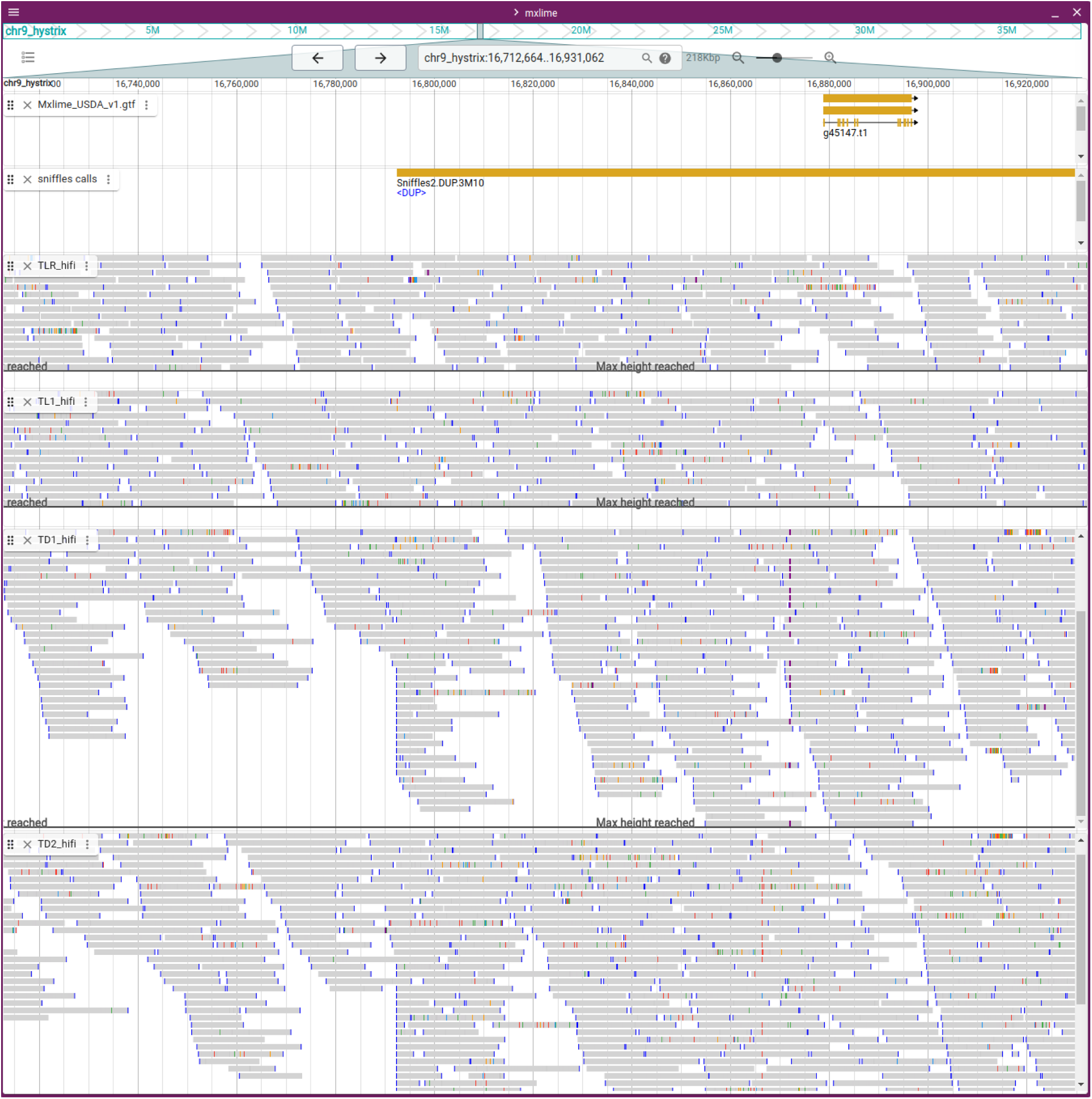
JBrowse2 view of large duplication event unique to thorned individuals.

**Figure S9.**
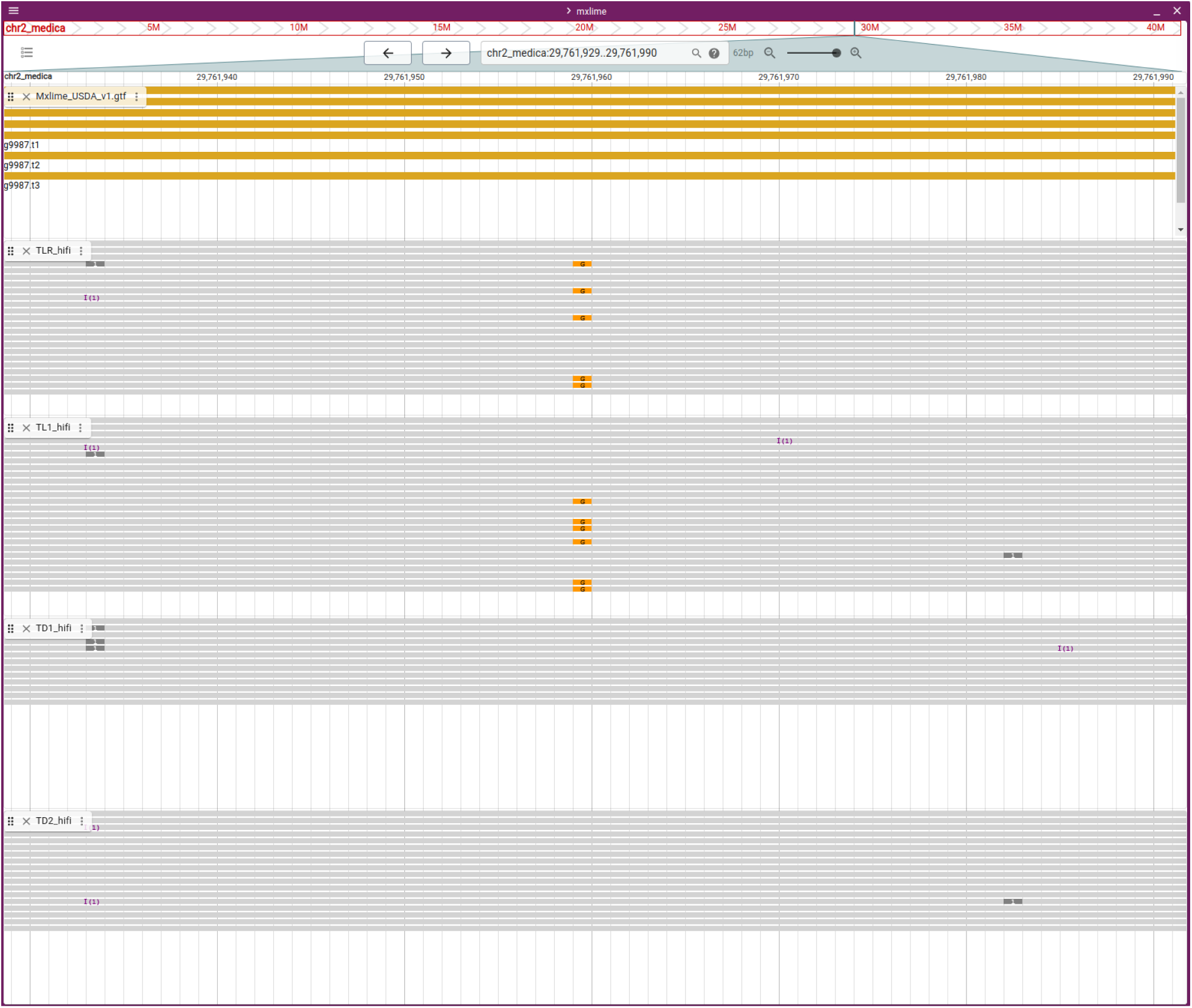
JBrowse2 view of a mosaic SNP in thornless individuals.

**Table S1.** Sequencing data used for this manuscript.

| Library | Read format | Raw data (Gb) | Number of reads | SRA |
| --- | --- | --- | --- | --- |
| Mxlime_HiFi | Circular consensus | 45.1 | 3,477,920 | SRR30099116 |
| Mxlime_HiC | Illumina PE151 | 35.4 | 117,487,668 | SRR30099114 |
| Mxlime_flower_mRNAseq | Illumina PE151 | 15.3 | 50,677,025 | SRR30099115 |
| Mxlime_shoot_apex_mRNAseq | Illumina PE151 | 15.2 | 50,382,236 | SRR30099111 |
| Mxlime_leaves_mRNAseq | Illumina PE151 | 18.1 | 59,810,818 | SRR30099113 |
| Mxlime_fruit_mRNAseq | Illumina PE151 | 18.4 | 61,238,269 | SRR30099112 |
| Mxlime_TD1_HiFi | Circular consensus | 15.8 | 991,202 | DRYAD |
| Mxlime_TD2_HiFi | Circular consensus | 16.0 | 985,638 | DRYAD |
| Mxlime_TL1_HiFi | Circular consensus | 24.0 | 1,489,941 | DRYAD |

**Table S2.** Presence of telomeres for each subgenome, “(AAACCCT)_n_”.

| Chromosome | 5' telomere | 3' telomere |
| --- | --- | --- |
| Chr1_hystrix | present | present |
| Chr2_hystrix | present | absent |
| Chr3_hystrix | present | present |
| Chr4_hystrix | present | absent |
| Chr5_hystrix | present | present |
| Chr6_hystrix | present | absent |
| Chr7_hystrix | absent | present |
| Chr8_hystrix | present | present |
| Chr9_hystrix | present | absent |
| Chr1_medica | present | present |
| Chr2_medica | absent | present |
| Chr3_medica | present | present |
| Chr4_medica | present | present |
| Chr5_medica | present | present |
| Chr6_medica | present | present |
| Chr7_medica | present | absent |
| Chr8_medica | absent | present |
| Chr9_medica | present | present |
| Scaffold_21 | absent | present |
| Scaffold_22 | absent | absent |
| Scaffold_23 | absent | absent |
| Scaffold_24 | present | absent |
| Scaffold_25 | present | absent |
| Scaffold_26 | absent | absent |
| Scaffold_27 | absent | absent |
| Scaffold_28 | absent | absent |

